# Data-driven models reveal mutant cell behaviors important for myxobacterial aggregation

**DOI:** 10.1101/2020.02.08.939462

**Authors:** Zhaoyang Zhang, Christopher R. Cotter, Zhe Lyu, Lawrence J. Shimkets, Oleg A. Igoshin

**Affiliations:** Department of Bioengineering and Center for Theoretical Biological Physics, Rice University, Houston, Texas, United States of America; Department of Microbiology, University of Georgia, Athens, Georgia, United States of America

## Abstract

Single mutations frequently alter several aspects of cell behavior but rarely reveal whether a particular statistically significant change is biologically significant. To determine which behavioral changes are most important for multicellular self-organization, we devised a new methodology using *Myxococcus xanthus* as a model system. During development, myxobacteria coordinate their movement to aggregate into spore-filled fruiting bodies. We investigate how aggregation is restored in two mutants, *csgA* and *pilC*, that cannot aggregate unless mixed with wild type (WT) cells. To this end, we use cell tracking to follow the movement of fluorescently labeled cells in combination with data-driven agent-based modeling. The results indicate that just like WT cells, both mutants bias their movement toward aggregates and reduce motility inside aggregates. However, several aspects of mutant behavior remain uncorrected by WT demonstrating that perfect recreation of WT behavior is unnecessary. In fact, synergies between errant behaviors can make aggregation robust.

## 1 Introduction

Development is one example of multiscale emergent behavior in which molecular interactions between cells allow self-organization into multicellular patterns. One of the most remarkable features of all types of development is how robust it is in the face of genetic and environmental perturbations, suggesting that backup systems are in place [33]. While molecular genetics has identified mutations that impede multicellular development, even single mutations create downstream effects that influence multiple aspects of cell behavior and physiology. It is frequently difficult to ascertain which of the behavioral changes are deleterious to development and which can be tolerated. Here we develop a new approach that leverages data-driven modeling to determine whether a statistically significant trend in cell behavior results in a biologically significant alteration of the multicellular program. We demonstrate this approach by focusing on the full or partial rescue of the mutants during the multicellular development of *Myxococcus xanthus* biofilms.

*Myxococcus xanthus* is a rod-shaped member of the delta-Protobacteria with a lifecycle centered around surface motility of cells in a biofilm. *M. xanthus* has evolved multiple social mechanisms such as S-motility [14] and C-signaling [6,22,32] to achieve coordinated group behaviors such as predation [31], rippling [4,15,34] and development [34,42]. Upon amino acid limitation, *M. xanthus* cells move into three-dimensional aggregates called fruiting bodies where they sporulate [19,24,29]. Recent studies based on cell tracking have provided unprecedented detail of cell movement during development [9]. In combination with mathematical modeling, these datasets unambiguously identified individual cell behaviors that are essential for aggregation [9,41]. These behaviors include reduced movement inside the aggregate and bias in the directed movement toward the aggregation centers, likely via chemotaxis [41]. This methodology provides an unprecedented window into developmental behavior that is presently difficult to realize in larger organisms with thicker tissues or longer cell migration routes, such as the vertebrate neural crest or in disease states such as tumor metastases.

In this work, we examined reciprocal interactions between WT cells mixed with non-developing mutants. More so than other bacteria, *M. xanthus* cell growth and development depend on neighboring cells, diffusing molecules, and the surrounding biotic and abiotic environment. To determine the factors that contribute to developmental robustness we employed conditional mutants that were unable to develop on their own but will develop when mixed with WT cells. It is expected that the mutants respond to at least some of the conditions established by WT cells in the field of developing cells. The extent of the response is expected to reveal signaling and sensory transduction pathways that are essential for WT development and are defective in the mutants.

The extent of WT rescue of two mutants is examined in this work. The first of these, contains a mutation in the *pilC* gene. PilC is an inner membrane protein located at the base of the pilus where it interacts with PilB and PilM to mediate pilus assembly [3,8]. This mutation interrupts pilus production [36] and consequently S-motility, one of the two motility systems in *M. xanthus* [26,28]. Aggregation can occur with the help of the A-motility system, which uses a novel molecular motor and focal adhesion complexes [12,27]. However, most S-system mutants fail to develop because they cannot produce an extracellular matrix (ECM) that is both essential for S-motility and vital for development. The ECM is required for some types of chemotaxis [20,21] as well as for cell cohesion, which could inhibit motility inside the aggregate [1,2]. As shown in this work, *pilC* mutants cannot aggregate on their own but marginally improve when mixed with wild-type cells. The second mutation is a deletion of the *csgA* gene, which inhibits the production of one or more intercellular signals that are required for aggregation and sporulation [13]. While CsgA signaling exerts control over most of development, the precise nature of the signals and their sensory pathways is only beginning to be revealed ([23]). *csgA* cells do not form fruiting bodies on their own [34], respond much more completely to a WT cell developmental field than *pilC* [25]. Although much is known about *M. xanthus* aggregation [25,34,35,40], few quantitative data sets describe mutant cell movement during aggregation and the mechanism of their rescue [18].

To identify motility behaviors affecting mutant cell aggregation, we extended our previously developed approach that combines individual cell tracking with simulations driven by the accumulated cell behavior data [9]. Directly applying experimental cell data to simulations allowed us to fully investigate the effect of each change in the mutant motility behavior on their aggregation. The results demonstrate that the WT developmental field is robust enough to nearly completely restore *csgA* development. By comparison, the *pilC* mutant has two striking sensory deficits that diminish its ability to accumulate inside the fruiting bodies. By exchanging particular aspects of cell behavior between WT and mutant cells, our agent-based modeling was able to pinpoint specific differences in cell behavior that are most biologically significant.

## 2 Results

### 2.1 Quantifying aggregation dynamics in mixtures of wild-type and mutant strains

Fluorescence microscopy was used to quantify the behavior of mutant cells at both single-cell and population levels. A small fraction of cells expressing the fluorescent protein tdTomato were mixed with cells expressing eYFP. Each cell expressing tdTomato is bright enough to be segmented and tracked, allowing quantification of their behaviors, whereas the weaker eYFP signal was used to quantify cell density during aggregate growth [9].

When either *pilC* or *csgA* cells are mixed with differentially labeled cells of their genotype, no aggregates were observed and the distribution of cells is nearly uniform at the final time-point, i.e. at T=5h (fig. 1AB). Application of the 2-D Kolmogorov-Smirnov test [30] to cell positions shows that the null-hypothesis of the uniform distribution of labeled cells cannot be rejected (p-value¿0.95). Conversely, when tdTomato labeled *csgA* cells are mixed with eYFP labeled wild-type (WT) cells, *csgA* cells are overrepresented in the aggregates (fig. 1D). The distribution of the cells is clearly non-uniform (p-value¡0.05). For *pilC* cells mixed with WT cells, the rescue is less pronounced (fig. 1C) and there is not sufficient evidence to reject the null hypothesis of uniform distribution of labeled cells (p-value=0.64). Below we describe a more sensitive metric to quantify aggregation rescue of mutant cells. As a comparison, fig. S1 shows the aggregation result of WT cells

**Figure 1.**
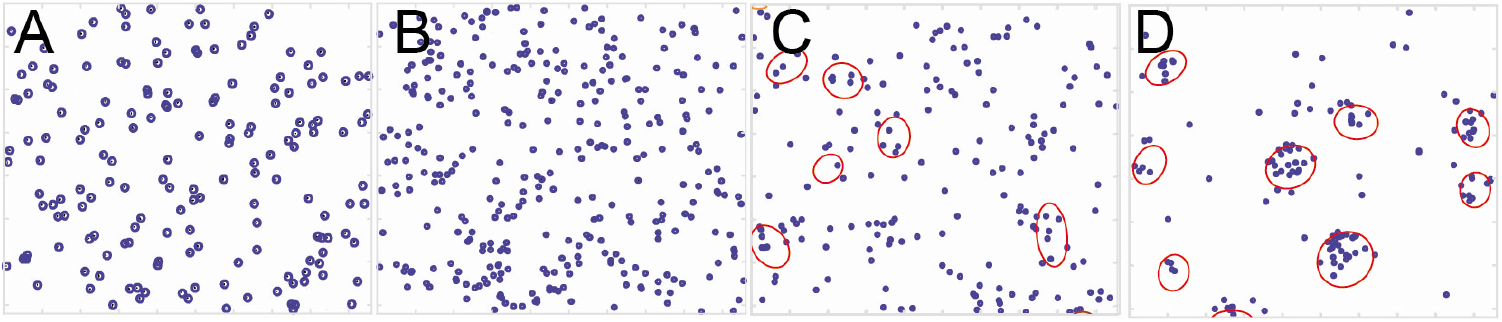
Cell distribution at the final frame of the experimental movies. Cells are segmented and shown as blue circles at the centroids of labeled cells. Red ellipsoids indicate the boundaries of aggregates segmented from the image after cells were filtered. (A) *pilC* cells alone. (B) *csgA* cells alone. (C) *pilC* mixed with WT cells. (D) *csgA* cells mixed with WT cells.

To quantify aggregate positions, densities, and sizes, we filtered out the tdTomato signal then used the eYFP intensity to estimate cell density. This data was used to segment the aggregates and detect their boundaries and positions. For segmentation of the images in which aggregation was observed (mutant strains mixed with a majority of WT cells), we determined a threshold intensity that separates aggregates from the background using K-means clustering on the light intensity of each pixel in the final frame of the experimental movies. Dividing the light intensity of pixels into two clusters gives the threshold of light intensity for aggregates. Applying the same threshold throughout the sequence of time-lapse imaging, we can compare aggregate growth for different experiments. To compare the aggregation rate across different sets of experiments, we use the average aggregate size fraction, *F_agg_*(*t*), i.e. the total area of aggregates in each frame corresponding to time (*t*) divided by the field of view area. The results (fig. 2A) indicate that aggregation of WT mixed with *pilC* cells is slightly slower than WT aggregation (dataset from [9]. On the other hand, WT cells mixed with *csgA* show faster aggregation. However, at the final time point, datasets lead to approximately the same area covered by aggregates, *F_agg_* (*t_final_*). Given that WT cells represent the overwhelming majority (¿99.9%) of the cells it is unlikely the observed differences are directly attributable to the presence of mutant cells. Instead, these differences are likely due to a slight variation of experimental conditions. Indeed, different biological repeats of the mixture experiments show differences in the aggregation dynamics (fig. 2BC). Therefore, previously used metrics to characterize aggregation such as the fraction of cells within the current area of aggregates could be overly sensitive to this variability.

**Figure 2.**
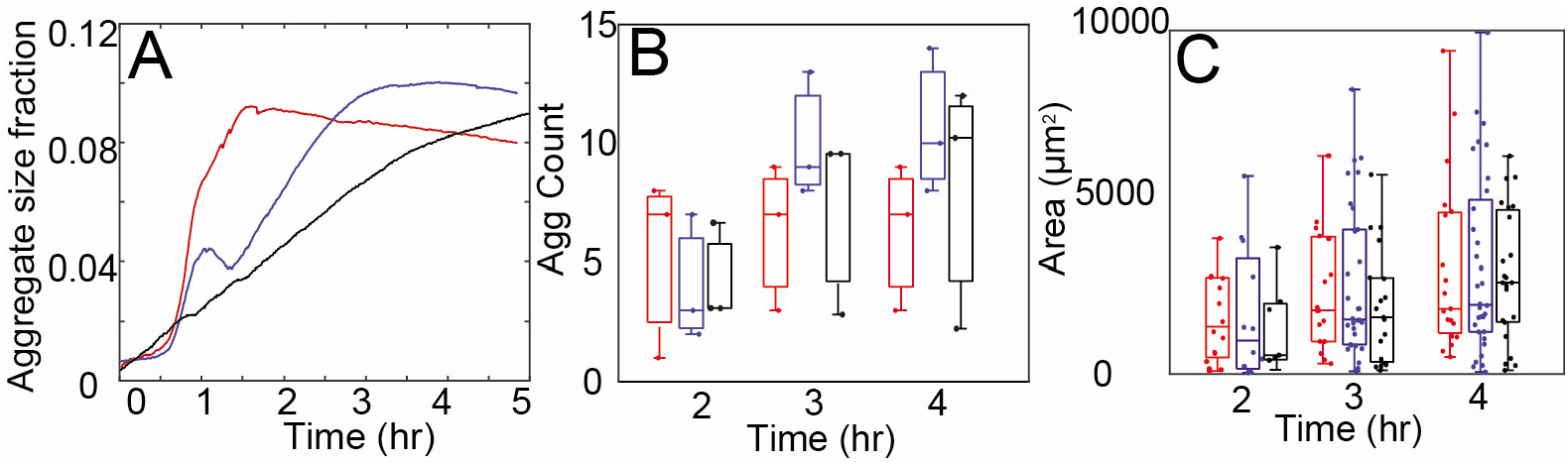
Aggregation variation between experiments: (A): WT cell aggregation rates vary between experiments. Y-axis is the aggregates area divided by the total area of the field of view. The red line is the average aggregation rate in experiments mixing *csgA* with WT cells. The blue line is the average aggregation rate of WT cells only [9]. The black line is the average aggregation rate in experiments mixing *pilC* with WT cells. (B, C): Aggregate numbers (B) and each aggregate area (C) in experiments. Red is *csgA*, blue is WT, and black is *pilC*. Horizontal lines inside the boxes indicate the distribution median. Tops and bottoms of each box indicate 75th (q3) and 25th (q1) percentiles, respectively. Dots in (B) are aggregate number in each experiment and dots in (C) are each aggregate area in experiments

To quantify the distribution of the tracked cells relative to the aggregates in a way that is robust to the variability of aggregation rate, we decided to focus on the fraction of cells accumulated inside the final-frame boundaries of the aggregates. If the tracked cells were uniformly distributed, we would expect that fraction to be equal to the fraction of area covered by aggregates, i.e. *F_agg_*(*t_final_*). Therefore, to see if labeled cells are overrepresented we focus on:

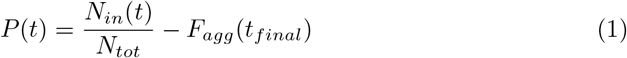

Here *N_in_* is the number of tracked cells inside the final aggregate area and *N_tot_* is the total number of tracked cells over the total field of view area. We do this calculation for each frame (at time *t*) and use it to quantify the aggregation rate of labeled cells.

The results for *P*(*t*) quantification for aggregation of *csgA* mixed with WT (red) and *pilC* mixed with WT (black) cells are shown fig. 3A. To compare it with WT only aggregation, we use a dataset of [9] to compute the same quantity (fig. 3A, blue line). The result shows that *csgA* has a similar aggregation rate to WT cells. In the final frame, the number of cells inside aggregates is larger by 50% compared with the total cell number, *P*(*t_final_*) ~ 0.5. In contrast, *pilC* cells show much weaker aggregation *P*(*t_final_*) ~ 0.1. To test if overrepresentation of *pilC* mutants inside the aggregate is statistically significant, we performed a z-test. The null hypothesis is that the *pilC* cells are randomly distributed, therefore the mean of *P*(*t_final_*) is 0. The p-value for accepting the null hypothesis is 0.002, indicating that the *pilC* mutant is partially rescued by WT cells.

**Figure 3.**
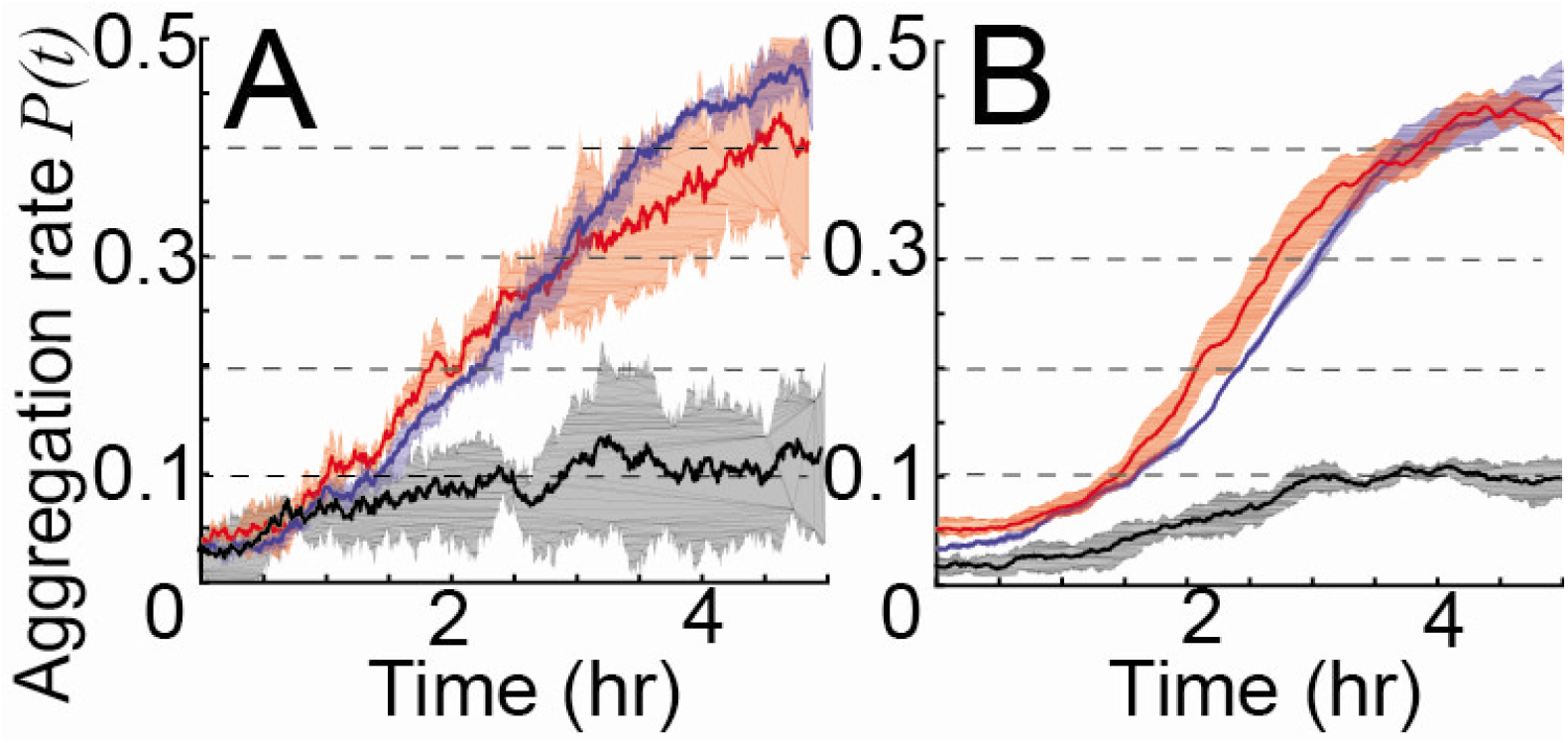
Comparison of aggregation rates (Eq. 1) between experiment (A) and simulation (B). Solid line is the average value and shaded area is a standard deviation for each time point. Red, blue and black colors correspond to WT,csgA mixed with WT and *pilC* mixed with WT respectively.

### 2.2 Motility behaviors of rescued *pilC* and *csgA* cells differ from WT cells

To quantify single-cell behaviors, the cell trajectories were discretized into segments using the same method as in [9]. The resulting segmented trajectories were then quantified as either persistent or non-persistent run vectors. Persistent runs are interpreted as cells moving along their major axis using one or both motility systems whereas as non-persistent runs correspond to “stops” (or pauses) in progressive movements, during which cells can perhaps be pushed around by other cells. A run vector begins at a change of state (persistent to either non-persistent or reversal) and ends at the next change of state. The properties of the resulting run vectors, such as duration (time between state changes) and speed (Euclidean distance over time) were used to quantify single cell behavior during aggregation. The run vectors were also labeled with the distance to the nearest aggregate boundary and moving direction relative to the nearest aggregate center. Previous work has shown that WT cells have longer run durations when running towards an aggregate (bias effect) and cells decrease their motility inside aggregates (”traffic-jam” effect) [9,16]. These effects have been shown to be important for aggregation [9, 35, 41]. To quantify traffic jam and bias effects, we focus on the relationship between run vector properties and their distance and direction relative to aggregates.

To study the relationship between the run vector properties and the distance to aggregates, we divided the run vectors into 2 groups: those inside aggregates and those outside. Then we calculated the mean duration and speed for the persistent and non-persistent state in each group (fig. 4). We find that both WT and mutant cells mixed with WT cells display a traffic-jam effect since they all have shorter persistent run durations and longer non-persistent run durations inside aggregates (fig. 4BD). To quantify the bias in run duration, we divided the run vectors into 2 groups: those running towards aggregates and those running away. Then we define the bias ratio by

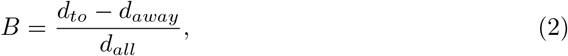

where *d_to_* is the average run duration of cells going towards aggregates, *d_away_* is the average run duration of cells going away from aggregates, and *d_all_* is the average run duration of all cells. fig. 4C shows that each mutant mixed with WT cells has a bias ratio greater than 0, though both are less than WT.

**Figure 4.**
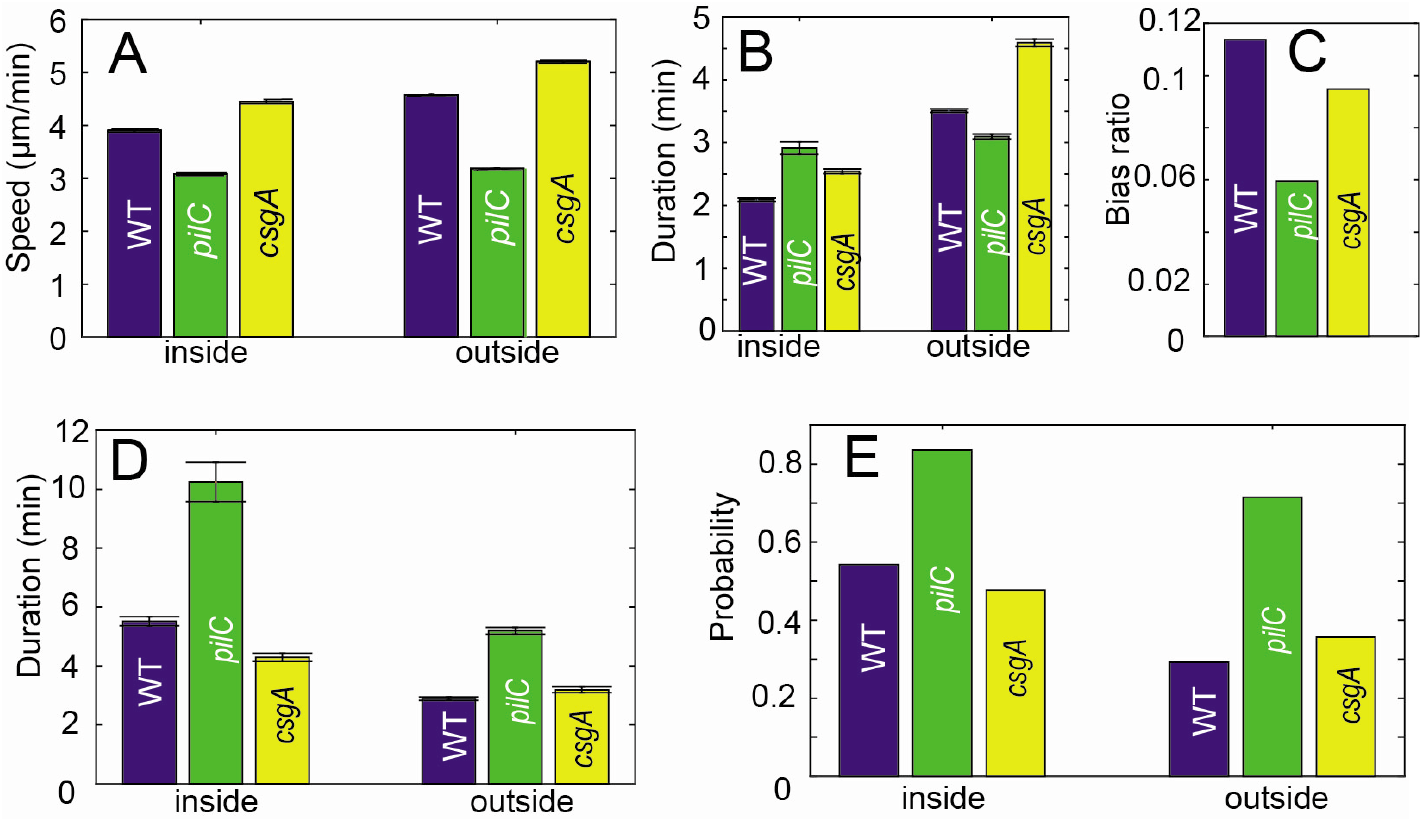
Experimental results of *pilC* and *csgA* cell behavior when mixed with WT cells compared with data from WT cells alone. Error bars represent bootstrapped 95% confidence interval of the means. (A): Persistent state speed of cells inside and outside aggregates. (B): Persistent state duration of cells inside and outside aggregates. (C) Bias ratio as defined in Eq.2 shows the tendency of cells to extend their runs when approaching the aggregates. (D): Non-persistent duration of cells inside and outside of aggregates. (E): Probability of transitioning into a non-persistent state after a persistent run for cells inside and outside aggregates.

To compare the traffic jam effect of *pilC* cells mixed with WT cells, we compared the speed and state durations of *pilC* and WT cells. In general, *pilC* cells exhibit longer stop durations, more frequent stops, and slower speeds suggesting that loss of S-motility has compromised their overall mobility. Unlike WT cells, *pilC* cells show less than a 5% speed reduction inside aggregates during the persistent state (fig. 4A) and only show 7% shorter persistent run durations (fig. 4B). Furthermore, *pilC* cells show less bias in their run duration (fig. 4C). As compared with WT and *csgA*, smaller differences between cell behaviors inside and outside aggregates may reduce the traffic jam effect, thereby impeding aggregation of *pilC* cells. On the other hand, *pilC* cells show a longer non-persistent duration and a higher probability of transitioning to the non-persistent state. However, the difference between the transitioning probability between inside and outside aggregates is smaller (fig. 4D, E).

Similarly, we compared the traffic-jam and bias effects of a *csgA*-WT mixture with WT cells. While *csgA* speed is ~ 20% faster than WT cells inside aggregates, *csgA* cells show proportional speed reduction inside aggregates (fig. 4A). Similar to WT cells, *csgA* cells also have shorter persistent durations inside aggregates (fig. 4B), and longer non-persistent durations (fig. 4D). Moreover, *csgA* cells increase their probability of transitioning to the non-persistent state when inside the aggregates. However, the difference of this probability between inside and outside the aggregates of *csgA* cells is smaller than that of WT cells (fig. 4E). All of the above behaviors reduce the motility of *csgA* cells inside the aggregates, likely creating a WT-like traffic-jam effect.

In comparison with *pilC* cells (fig. 4), *csgA* cells likely have a stronger traffic jam effect due to a more pronounced reduction in speed and persistent run duration inside the aggregates. On the other hand, their traffic-jam effect is expected to be weaker than WT due to reduced differences in non-persistent duration and probability between inside and outside. The *csgA* cells also have a weaker bias than WT cells (fig. 4C). It remains to be seen why, despite a somewhat weaker bias and traffic jam effect, about the same proportion of *csgA* cells accumulate in the aggregate as WT cells (fig. 3).

### 2.3 Data-driven models can match the aggregation dynamics of *pilC* and *csgA* cells based on the quantified motility parameters and their correlations

To more stringently test the effect of cell behaviors on aggregation, we extended the data-driven model approach used in our previous work [9] to model experiments with mixtures of two strains. To this end, we introduce a population of two agents corresponding to WT and mutant (either *pilC* or *csgA*) cells. Agent behaviors are chosen from the experimental data using K-nearest neighbor (KNN) sampling based on simulation time and the agent’s distance and moving direction relative to the nearest aggregate. Given that the overwhelming majority of cells in the experiments are WT, we only use WT agent density to detect aggregates. This way, WT agents affect the behavior of mutant agents but not *vice versa*. At each time step, the WT density profile is estimated from the WT agent positions by kernel density estimation (KDE) [5] and the aggregates were then detected from the density profile. Thereafter, we pick agent behaviors and move agents accordingly. Each simulation was run for 5 hours, after which we calculated the aggregation rate *P* (*t*) as we did for the experiment. Simulations containing *csgA* agents mixed with WT agents display an aggregation rate similar to that of WT agents, whereas simulations with *pilC* agents exhibit much weaker aggregation (fig. 3B). Comparing the results of these simulations to the experimental measurements (fig. 3A), we concluded that the model can reproduce the aggregation dynamics for WT and each mutant cell mixture with WT. In other words, dependencies (correlations) included in the sampling of agent behavior contain sufficient information to recapture observed aggregation dynamics.

As a control for the previous simulations, we performed simulations where we removed all dependencies such that agent behavior is randomly chosen from the whole data set. As expected, we did not see any aggregation for mutant mixtures or WT agents (fig. S2 A-C). This result shows that some combination of cell behavior dependence on time, distance, and direction to nearest aggregate is essential for aggregation. Since there are many cell behavior dependencies in this model, our next step is to find which dependencies are more important for aggregation.

Previous work on WT aggregation has shown that cell behaviors are different at different times during development and this time-dependence of cell behaviors affects aggregation dynamics [9]. To determine whether time dependence is important for mutant cell aggregation, we performed simulations where agent behavior does not depend on time. Removing time-dependence for WT aggregation causes *P*(*t_final_*) to drop from ~ 0.45 to ~ 0.35 (fig. S2F), which confirms our previous result [9] that time dependence helps WT aggregation. However, removing time dependence for mutant agents (while keeping it for WT agents), does not affect aggregation dynamics for either *pilC* (fig. S2D) or *csgA* (fig. S2E). This shows that behavior dependence on time is not important for mutant cell aggregation.

To further illuminate differences between WT, *cgsA* and *pilC* strains, we used our simulations to quantify the fraction of cells that enter and exit aggregates as a function of time (fig. 5). The results indicate notable differences. Comparing WT and *cgsA*, we can see that, although a more *cgsA* cells reach the aggregates (red line in fig. 5A vs B), a larger fraction of these cells leave(green line in fig. 5 A vs B). Therefore, we can hypothesize that reduced ”traffic-jamming” of *cgsA* cells is compensated by increase in motility. On the other hand, for *pilC* cells, the motility defects make them less likely to reach aggregates and slightly less likely to stay, leading to only weak aggregation. In what follows we aim to text these hypothesis and relate these observations with the trends in cell behaviors quantified in (fig. 4).

**Figure 5.**
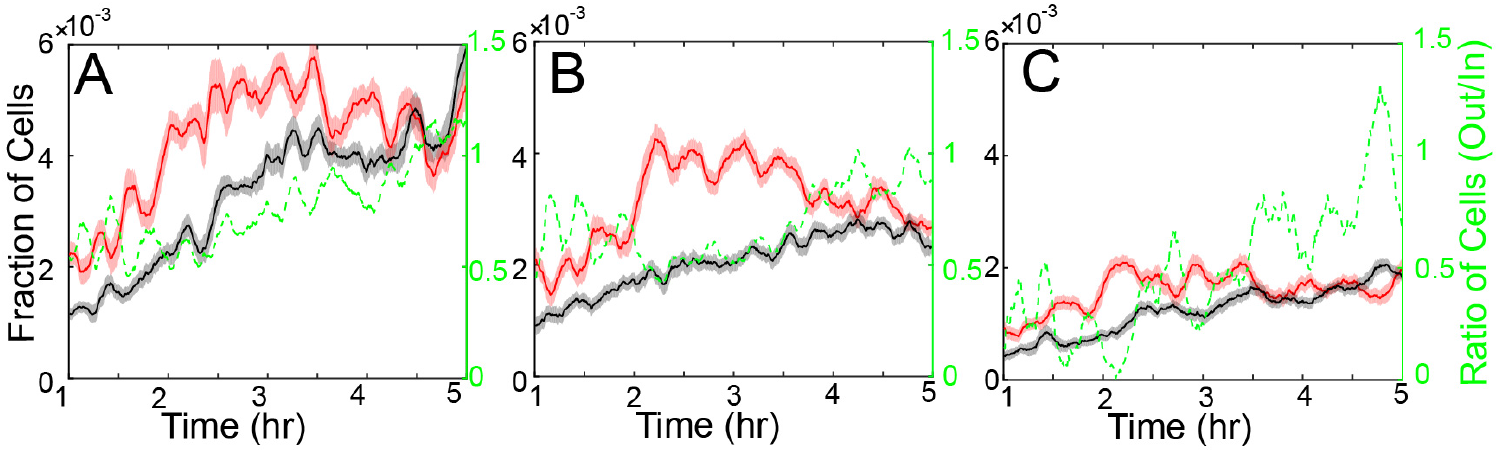
Analysis of agents moving in and out of aggregates at each time step for *csgA* (A), WT (B), and *pilC* (C). Red lines are the 10 min moving average of the fraction of agents moving into all aggregates at a given time. The fraction is calculated as a ratio of agents moving into the aggregate and the total number of agents simulated. Similarly, black lines are the 10 min moving average of the fraction of agents moving out of the aggregates. The shaded areas are 95% confidence interval of the moving mean. The right axis/dashed green show the ratio of number of agents moving out of aggregates divided by the number of agents moving in the aggregates.

### 2.4 Transitioning to and staying in the non-persistent state in aggregates does not help *pilC* and *csgA* aggregation

Given that the increase of non-persistent state duration increases the time cells spend inside aggregates, we hypothesized that this effect is an essential component of the traffic-jam effect and aids the aggregation of mutant cells. To test this hypothesis, we performed simulations where the non-persistent state duration for agents is not conditional on their position relative to the aggregate. Surprisingly, removing this dependence does not have an obvious effect on *pilC* (fig. 6A) or *csgA* (fig. 6B) and only leads to a modest decrease in WT aggregation (~ 0.05 or ~ 10% drop in *P(t_final_*); fig. 6C). This result shows that longer “stops” inside aggregates is not the main reason for successful aggregation.

**Figure 6.**
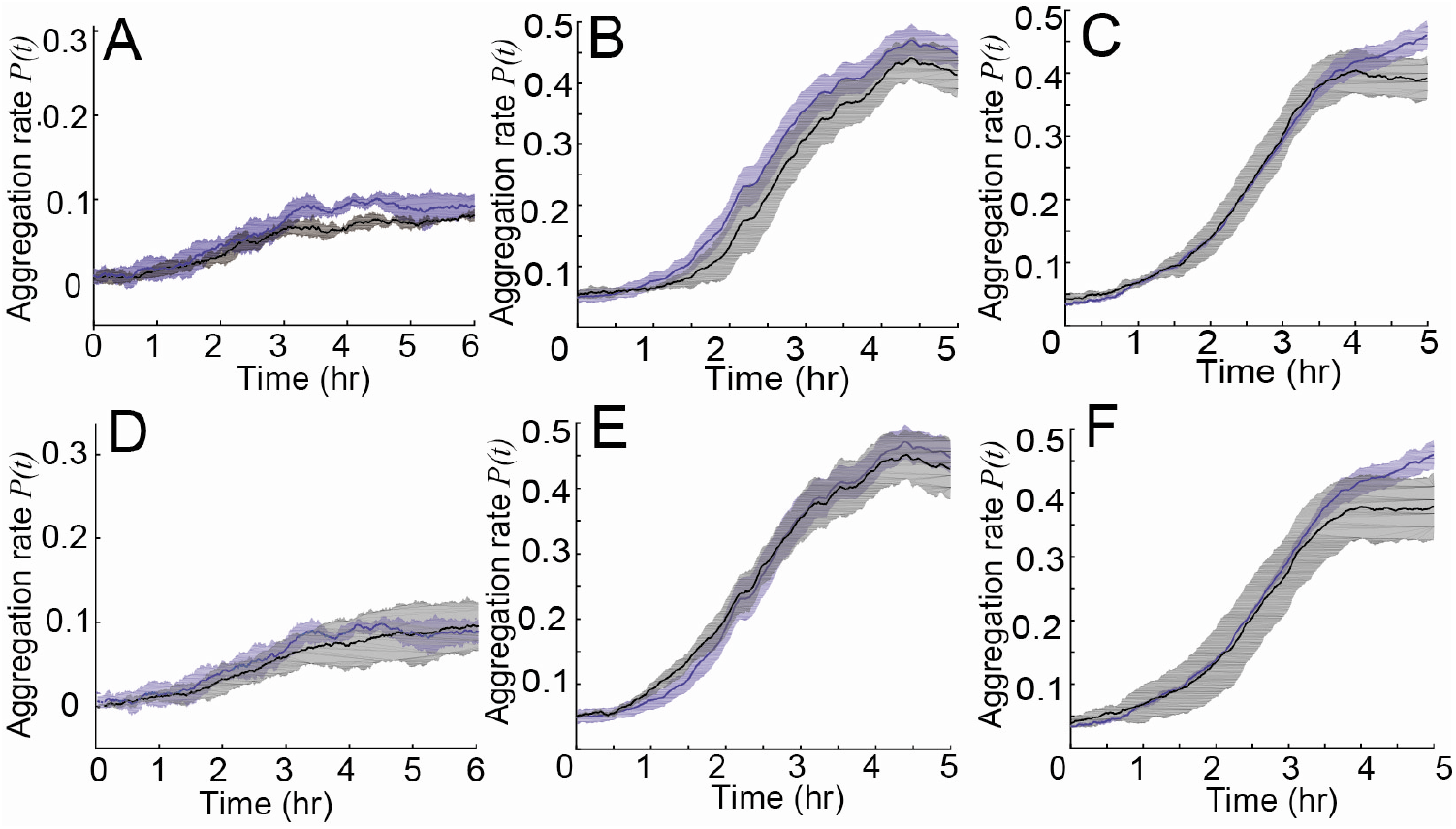
Comparison of the aggregation rate (quantified as *P*(*t*), Eq. 1) on y-axis) from simulations of *pilC* (A,D) and *csgA* (B,E) cells mixed with WT cells, and WT (C,F) cells alone. Blues line and shaded area is the simulation result under conditions where agent behavior is chosen from experimental data. Black lines represent simulations where non-persistent behavior does not depend on distance to aggregates (A-C) or in which probability to non-persistent state does not depend on distance to aggregates (D-F). Shaded areas show standard deviations.

To assess the effects of a higher probability of “stops” (i.e. non-persistent runs) inside the aggregates, we performed simulations where the probability of transitioning to a non-persistent state is independent of the agent’s position, (i.e. sampled from the same distribution inside and outside an aggregate). We discovered that removing this dependence does not affect aggregation for *pilC* (fig. 6D) or *csgA* (fig. 6E) and leads to only a ~ 0.07 (~ 15%) drop in *P*(*t_final_*) for WT (fig. 6F). It appears that longer non-persistent state durations and a higher probability of transitioning to the non-persistent state are not the main reasons for cell accumulation in aggregates. In summary, the difference in stopping probability and duration between inside and outside the aggregates are not critical for the traffic jam effect or can be compensated by other mechanisms.

### 2.5 Behaviors in the persistent state are critical for the aggregation

To test which persistent state behaviors are important for aggregation, we first removed the bias towards aggregates, which is the dependence of run duration on the angle between the moving cell and the closest aggregate. This leads to a ~ 0.03 drop in *P*(*t_final_*) for *pilC* (fig. 7A). For *csgA* (fig. 7B) and WT (fig. 7C), *P*(*t_final_*) drops 0.15~ 0.2. This result shows that bias in run duration is essential, more so for *csgA* and WT aggregation than *pilC* aggregation. This also agrees with fig. 4C where we showed that *csgA* and WT cells have larger bias ratios than *pilC* cells. However, given the overall poor aggregation of *pilC*, the decrease associated with lack of bias is still important and in relative terms is just slightly weaker than that of the other strains (30% reduction of final *P*(*t_final_*) for *pilC* vs 40% for *csgA* and 45% for WT).

**Figure 7.**
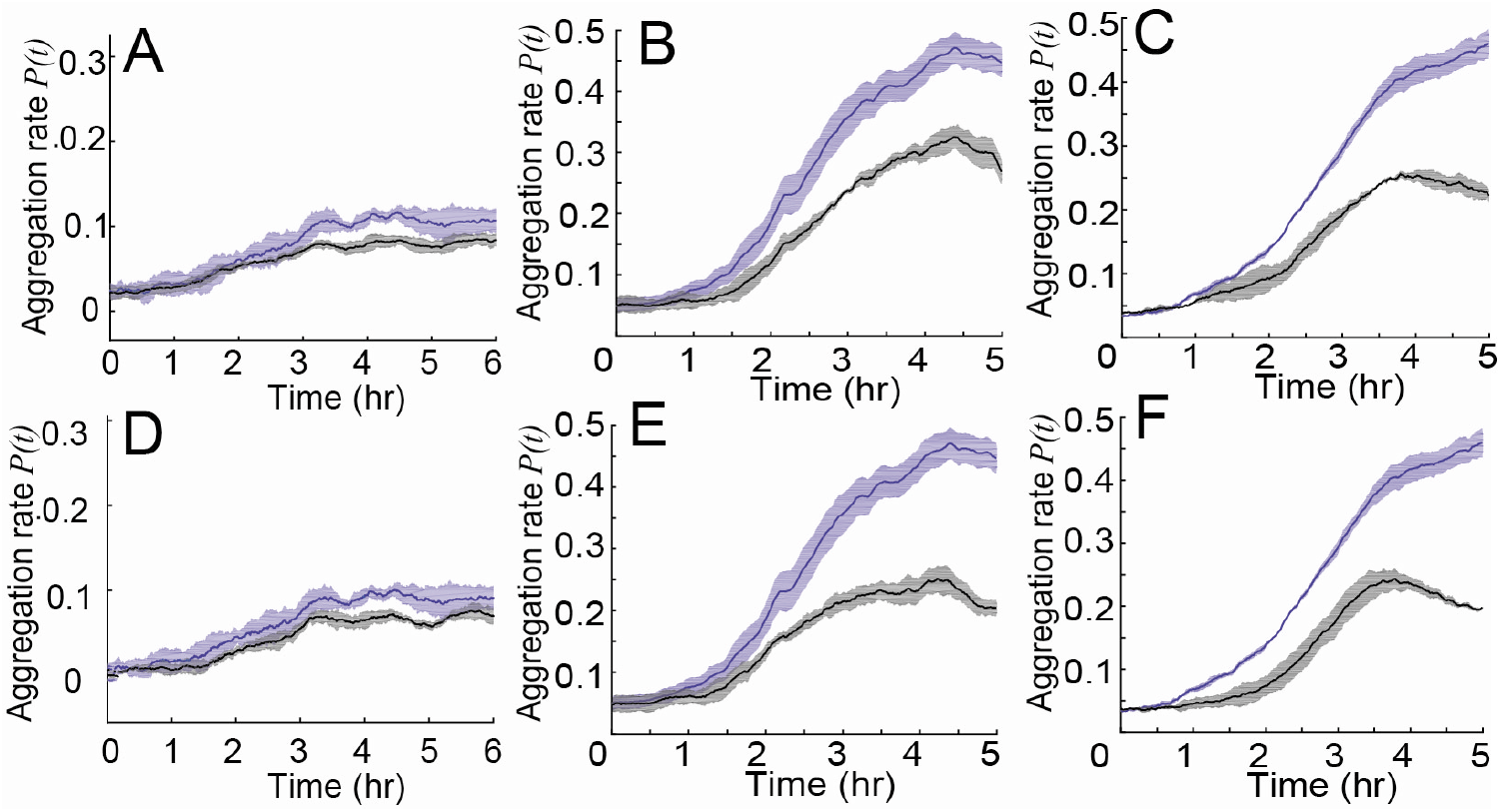
Comparison of the aggregation rate (quantified as *P*(*t*), Eq. 1) on y-axis) of simulated of *pilC* (A,D) and *csgA* (B,E) cells mixed with WT cells, and WT (C,F) cells alone. The blue lines and shaded area are simulation results under normal conditions. The black lines represent simulations where persistent behavior does not depend on run direction, i.e., without bias (A-C) or in which run duration does not depend on the distance to aggregates, i.e., without traffic jam (D-F). Shaded areas show standard deviations.

Next, we attempt to make the cells behave the same way inside and outside the aggregate to remove the traffic jam effect but maintain the bias. First, we removed persistent state speed and duration dependence on the agents’ distance to the nearest aggregate while keeping the dependence of run duration on the angle between the moving cell and the closest aggregate. The results show that removing the distance dependence decreases aggregation for all types of cells: *P*(*t_final_*) drops ~ 0.03 for *pilC* (fig. 7D) and drops ~ 0.25 for *csgA* (fig. 7E) and WT (fig. 7F) cells. Therefore, the reduction of speed and duration inside aggregates is important for aggregation. Interestingly, the reduction of speed and duration can also be considered a traffic-jam effect. Comparing the traffic-jam effects in the non-persistent state, i.e., longer duration and higher probability of non-persistent state inside aggregates, traffic-jam effects in the persistent state appear to be more important. Notably, removing persistent speed and duration dependence on distance decreases aggregation more in *csgA* and WT cells than in *pilC* cells. Even considering the poor aggregation of *pilC*, the relative decrease in aggregation is still weaker for *pilC* (30% reduction of final *P*(*t_final_*) for *pilC* vs 55% for *csgA* and 55% for WT). This shows that *csgA* and WT cells have a stronger traffic-jam effect than *pilC*, in agreement with fig. 4A and 4B.

### 2.6 Different motility behaviors of *pilC* and *csgA* cells explains the partial rescue of *pilC* and full rescue of *csgA*

The results thus far match the observed behaviors of mutant cells with their observed aggregation dynamics. Next, we try to determine which mutant cell behaviors are responsible for the different aggregation rates as compared with WT. To this end, we introduce a new “hybrid” simulation technique in which certain aspects of mutant and WT agent behaviors are swapped with one another or scaled to match the mean of another. For example, the experimental data shows that *pilC* mutants switch to the non-persistent state more frequently and stay in the non-persistent state longer (fig. 4D, E). To determine whether these behaviors contribute to weaker aggregation, we performed simulations where we swap some of the *pilC* motility behaviors with WT behaviors (fig. 8). When agents using the *pilC* probability of transitioning to the non-persistent state and WT data for other behaviors, aggregation drops (*P*(*t_final_*) drops ~ 0.2) (fig. 8A), but agents using WT probability of transitioning to the non-persistent state with *pilC* data for other behaviors does not improve *pilC* aggregation (fig. 8B). To further confirm that the decrease in aggregation is due to longer stops or a higher stopping frequency rather than some other feature of the *pilC* data, we performed simulations of WT cells where we only increased the non-persistent duration or non-persistent probability to match the average data of *pilC* cells (fig. S3). The aggregation rate is slowed compared to WT aggregation suggesting that frequent stops are one of the major impediments to *pilC* aggregation.

**Figure 8.**
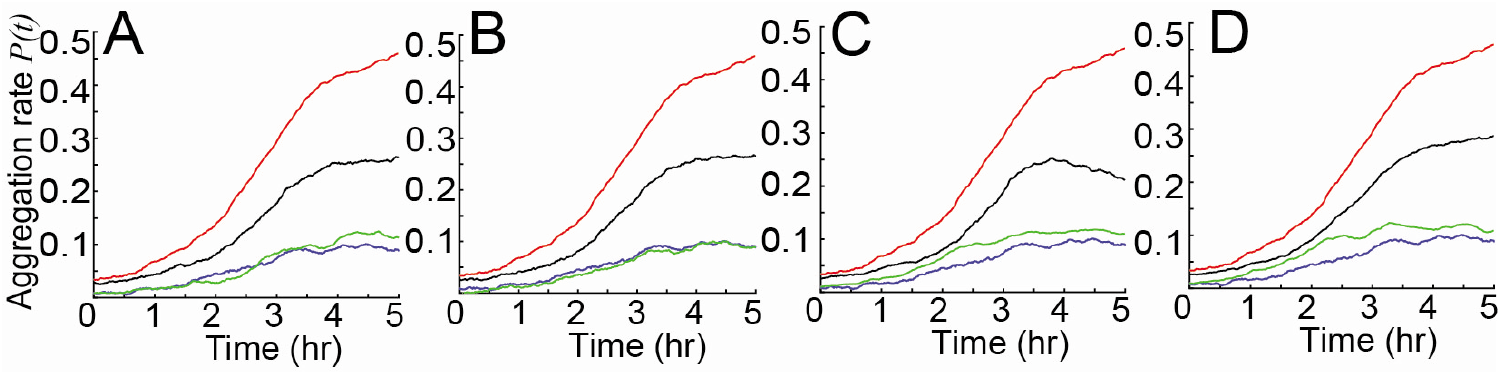
Simulations swapping WT data and *pilC* cell data demonstrate which mutant cell behaviors are sufficiently different from wild-type to affect the aggregation rate (quantified as *P*(*t*), Eq. 1) on the y-axis). Blue lines are simulation results of agents using *pilC* cell data. Red lines are simulation results of agents using WT cell data. Green lines are simulations of agents using *pilC* data with partial WT cell data. Black lines are simulations of agents using WT data with partial *pilC* cell data. (A): Green is agents using WT cell probability to non-persistent state and other *pilC* data. Black is agents using *pilC* probability to non-persistent state and other WT data. (B): Green is agents using WT cell non-persistent state duration and other *pilC* data. Black is agents using *pilC* cell non-persistent state duration and other WT data. (C): Green is agents using WT cell persistent state duration and other *pilC* data. Black is agents using *pilC* cell persistent state duration and other WT data. (D): Green is agents using WT cell persistent state speed and other *pilC* data. Black is agents using *pilC* cell persistent state speed and other WT data. Only mean values are plotted for clarity.

To learn how *pilC* persistent behaviors affect aggregation, we performed simulations where agents use the WT persistent duration data combined with other *pilC* cell data and vice versa (fig. 8C). Agents using *pilC* persistent duration combined with other WT data have reduced aggregation compared with WT (*P*(*t_final_*) drops ~ 0.25). Agents using WT persistent duration combined with other *pilC* data show improved aggregation over *pilC* (*P*(*t_final_*) increases ~ 0.02). This is not surprising since WT cells have a much stronger persistent duration bias and stronger bias leads to more complete aggregation. Finally, agents using *pilC* persistent speed combined with other WT data have reduced aggregation compared with WT (*P*(*t_final_*) drops ~ 0.2) whereas WT persistent speed combined with other *pilC* data improves *pilC* aggregation (*P*(*t_final_*) increases ~ 0.02) (fig. 8D). This is because *pilC* cells have similar speeds inside and outside aggregates whereas WT cells have slower speeds inside aggregates and this slowdown improves aggregation. Overall our results show that weak aggregation of *pilC* is due to slow speed, longer non-persistent durations, and a higher probability of transitioning to the non-persistent state.

For *csgA* mutants, fig. 4E shows that the difference for stopping probabilities and durations inside and outside aggregates is less pronounced than WT. To test whether these behaviors decrease aggregation, we performed a simulation where agents use WT data for probability of transitioning into the non-persistent state and *csgA* data for other behaviors. This simulation does not improve *csgA* aggregation (fig. 9A). But agents using *csgA* probability to transition to the non-persistent state and WT data for other behaviors cause *P*(*t_final_*) to drop ~ 0.05 compared with WT aggregation. Moreover, in fig. 9B, agents using *csgA* non-persistent duration and WT data for other behaviors show a slight decrease in aggregation compared with WT aggregation (*P*(*t_final_*) drops ~ 0.05). On the other hand, agents using WT non-persistent duration and *csgA* data for other behaviors show a slight increase in aggregation compared with *csgA* aggregation (*P*(*t_final_*) increases ~ 0.02). These results show that the differences in non-persistent state switching and duration between WT and *csgA* do not affect aggregation much.

**Figure 9.**
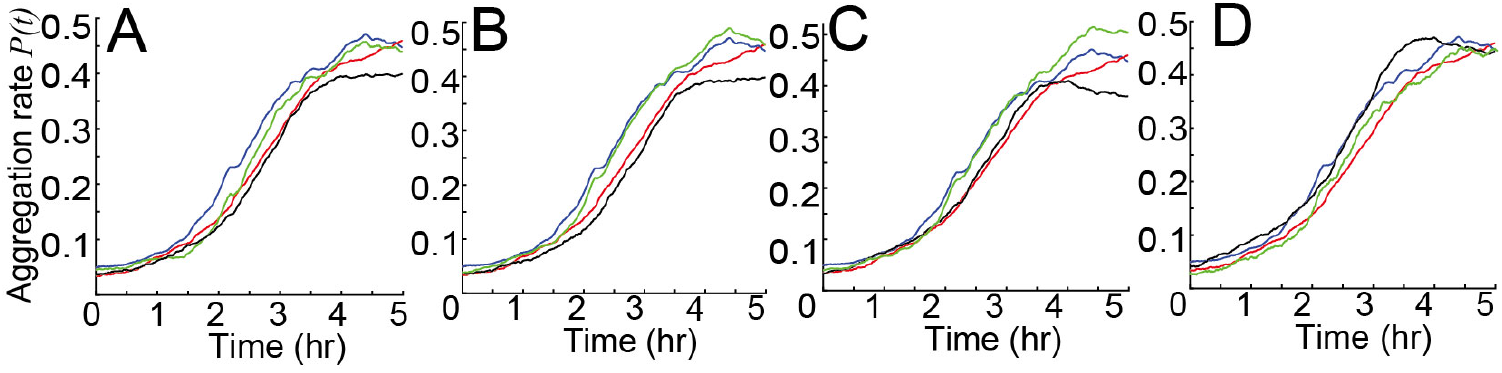
Simulations swapping WT data and *csgA* cell data demonstrate which mutant cell behaviors are sufficiently different from wild-type to affect the aggregation rate (quantified as *P*(*t*), Eq. 1) on y-axis). Blue lines are simulation results of agents using *csgA* cell data. Red lines are simulation results of agents using WT cell data. Green lines are simulations of agents using *csgA* data with partial WT cell data. Black lines are simulations of agents using WT data with partial *csgA* cell data. (A): Green is agents using WT cell probability to non-persistent state and other *csgA* data. Black is agents using *csgA* probability to non-persistent state and other WT data. (B): Green is agents using WT cell non-persistent state duration and other *csgA* data. Black is agents using *csgA* cell non-persistent state duration and other WT data. (C): Green is agents using WT cell persistent state duration and other *csgA* data. Black is agents using *csgA* cell persistent state duration and other WT data. (D): Green is gents using WT cell persistent state speed and other *csgA* data. Black is agents using *csgA* cell persistent state speed and other WT data. Only mean values are plotted for clarity.

To learn how *csgA* persistent behaviors affect aggregation, we performed a simulation where agents use persistent duration of WT cells and other behaviors of *csgA* cells (fig. 9C). This leads to a slightly better aggregation compared with *csgA* cells (*P*(*t_final_*) increases ~ 0.05). Agents using *csgA* persistent duration and other WT cell behavior show a slightly lower aggregation compared with WT cells (*P*(*t_final_*) drops ~ 0.07). Note that WT persistent duration has a bigger bias but shorter duration. To learn whether a shorter duration will decrease *csgA* aggregation, we performed a simulation where agents use *csgA* data but scale the persistent duration to match the average duration of WT cells. This led to a lower aggregation (fig. S4). These results show that the *csgA* weaker bias is partially compensated by the longer persistent duration.

Finally, to find whether the faster speed of *csgA* cells in the persistent state helps aggregation, we performed simulations where agents use *csgA* persistent speed and other WT behaviors(fig. 9D). This leads to faster aggregation, but *P*(*t_final_*) remains the same compared with WT. Moreover, agents using WT persistent speed and other *csgA* behaviors have a slightly slower aggregation rate compared with *csgA*. This result shows that *csgA* cells’ faster speed compensates for the weaker (compared with WT cells) bias in persistent duration and explains why *csgA* and WT cells show similar aggregation rates. Therefore, the rescue of collective behavior can occur even without the complete rescue of the underlying single-cell behaviors.

## 3 Discussion

In this work, we developed a novel methodology to assess which aspects of individual cell behavior are responsible for observed trends in collective self-organization. We show that this approach is both robust and versatile by examining how aggregation is restored when *csgA* and *pilC* mutants were mixed with wild type (WT) cells. The *csgA* rescue is quantitatively complete based on the percentage of cells in the aggregate whereas *pilC* rescue is marginal. As with our previous analysis of wild-type aggregation dynamics, we conclude that three features of cell behavior contribute to efficient accumulation in aggregates. First, the cells follow aligned paths that precede the appearance of aggregates such that their orientations are correlated with one another and with the direction to the nearest aggregate likely to appear along their path. That essentially reduces the search for aggregates to 1D. Second, due to the bias in persistent run durations, cells move longer when approaching an aggregate than when moving in the opposite direction. This biased random walk results in an increase in cell flux toward the aggregates and accelerate the aggregation dynamics. The biased walk appears to be due to chemotaxis [41] and several lipids are known to be chemoattractants [21]. Finally, once in aggregates, the cells are less likely to leave. Notably, all three strains displayed these three features, but their contributions are roughly proportional to the extent of aggregation with *pilC* being the worst.

Our approach enables a detailed examination of motility parameters that mediate a particular aggregation feature. For example, we extended the data-driven modeling approach for hybrid populations of agents that correspond to wild-type and mutant behaviors by using swapped-dataset sampling (table 1) to pinpoint cell movement features that are responsible for trapping cells in an aggregate. When cells enter an aggregate their speed decreases, the time spent in a persistent state decreases, their probability to transition to a non-persistent state increases, and the time spent in the non-persistent state increases. By swapping each of these features between WT and mutant in agent-based models we discovered that longer non-persistent state duration and a higher probability of transitioning to the non-persistent state are not the main reasons cells accumulate in aggregates. Rather, reduction in speed and run duration are the most critical features that trap cells in aggregates. It is possible that this result will inform those now searching for the enigmatic chemical signal that traps cells in an aggregate.

**Table 1.** Summary of performed simulation and effect of mutant behaviors on aggregation

| Observation | Simulation performed | Effect on aggregation |
| --- | --- | --- |
| *pilC* mutants switch to non-persistent state more frequently | *pilC* agents use WT data for the probability of transitioning to the non-persistent state and vice versa | Frequent stops slow-down aggregation (fig. 8A), but the final aggregation result is similar after a longer simulation time (fig. S3) |
| *pilC* mutants stay in non-persistent state longer | *pilC* agents use WT data for the duration and speed of the non-persistent state and vice versa | Longer stops slow-down aggregation (fig. 8B), but the final aggregation result is similar after a longer simulation time (fig. S3) |
| Run duration for *pilC* mutants shows lesser dependence on cell density and smaller bias | *pilC* agents use WT data for persistent duration and vice versa. | Smaller bias and difference between inside and outside aggregates for run durations impedes aggregation (fig. 8C). |
| *pilC* mutants do not show speed reduction inside the aggregates. | *pilC* agents scaled persistent speed to match WT data for and vice versa. | Lack of speed reduction inside the aggregates impedes aggregation (fig. 8D). |
| Stopping probabilities inside and outside aggregates is less pronounced for *csgA* mutants | *csgA* agents use WT data for the probability of transitioning to the non-persistent state for and vice versa | The density-dependence of stopping probability does not have a major effect on aggregation (fig. 9A). |
| Stop durations inside and outside aggregates is less pronounced for *csgA* mutants | *csgA* agents use WT data for the duration and speed of the non-persistent state and vice versa | Density-dependence of stopping slightly impedes aggregation (fig. 9B). |
| *csgA* mutants have a weaker bias but longer duration in the persistent state compared with WT | *csgA* agents use WT data for the duration of the persistent state<br>Scaled the persistent duration of *csgA* agents to match mean values of WT | While longer persistent duration helps *csgA* aggregation (fig. S4), the weaker bias has an opposite and stronger effect. Overall, compared with other behaviors, *csgA* persistent duration impedes aggregation more (fig. 9C). |
| *csgA* mutants have faster speed in persistent state | *csgA* agents use WT data for the persistent speed and vice versa | Faster speed speeds up aggregation (fig. 9D). |

Our approach also revealed that *csgA* achieved complete rescue despite differing in most WT motility parameters. Most notably, *csgA* cells show longer persistent run distances resulting from faster speeds and longer durations in the persistent state, compensate for a reduced biased random walk to produce similar aggregation to WT cells. Genetic studies have shown that *csgA* cells possess both motility systems. The mutant fails to develop specifically because it fails to produce one or more essential developmental signals that mediate aggregation in addition to a signal that induces sporulation. While the mutant clearly responds to the wild-type signal(s), it would appear that the *csgA* mutation causes a slight downstream effect in perception or motility regulation that reduces the bias. The chemical nature of the aggregation signal(s) remains unknown and but seems to be a development-specific lipid-like molecule [6]. The signal is unlikely to be the EPS used in S-motility as *csgA* produces normal levels of EPS, shows normal agglutination, and possesses S-motility.

Finally, we attempted to learn why *pilC* rescue is so poor. *pilC* cells are significantly attenuated for all behaviors that are important for wild-type aggregation. The swapped dataset sampling approach revealed that the failure of *pilC* cannot be attributed to one or two specific changes. *pilC* cells lack pili and EPS. S-motile cells use the pilus to attach to EPS on adjacent cells, and retraction of a motor at the base of the pilus pulls the cell forward. *pilC* cells have significantly decreased bias which our results suggest is due to reduced speed, reduced run durations, and increased frequency of transiting to the non-persistent state. As the WT cells would be expected to provide normal levels of EPS and other required signals, *pilC* cells clearly lack the appropriate response. This is a striking finding in view of the observation that neither pili nor S-motility are required for aggregation. Some S mutants, like *pilA* (which encodes the pilus structural protein) and *pilT* (which encodes the pilus retraction motor), can aggregate using only the A-motility system [10,37]. The results point to a downstream defect, perhaps related to the perception of lipid chemoattractants, which has been noted in certain S mutants but not examined specifically in *pilC*. The *pilC* mutant also has a diminished traffic jam effect. *pilC* cells frequently leave aggregates and ultimately are about 4-fold less abundant than *csgA* or WT cells. Again, the as yet unknown signal(s) used to hold cells in aggregates should be in sufficient concentration leading one to suspect that the problem is more specifically due to *pilC* response to the signal.

There is also a striking difference in the rescue of the two mutants for sporulation by WT cells. As a benchmark, WT cells form 0.14 spores per input cell with the remaining cells undergoing alternate developmental fates such as programmed cell death and formation of peripheral rods. WT cells efficiently rescue the sporulation of *csgA* mutants. *csgA* cells alone form 2 x 10^−5^ spores per input cell which increases to 0.15 spores per input cell in the presence of WT cells. In contrast, WT cells do not rescue *pilC* sporulation. *pilC* cells alone form 3 x10^−3^ spores per input cell which increases minimally to 4 x 10^−3^ spores per input cell in the presence of WT cells. Sporulation is thought to have little bearing on aggregation since it occurs after aggregation is complete. Nevertheless, these results clearly support the ideas developed in this work for aggregation that *csgA* cells are proficient in responding to signals unlike *pilC*.

Data-driven modeling approaches can identify important correlations that drive self-organization as a first step to understand the mechanisms. For example, in our previous work we have combined data-driven and mechanistic agent-based modeling to show that steric alignment and slime-trail following are sufficient to explain how cells align during aggregation [41]. Moreover, we also showed that chemotaxis rather than contact-based signaling is responsible for bias in reversal times for cells moving towards vs away from the aggregates [41]. However, the the exact molecular pathway(s) responsible for chemotaxis have yet to be determined. Furthermore, molecular mechanisms for the interactions that decrease the residence time of cells inside the aggregates (traffic-jam effects) are not clear. Another interesting observation is that both mutant strains considered here displayed differences from wild-type behavior in all the behaviours driving aggregation. That fact can indicate complex or possibly pleiotropic interactions between the pathways controlling cell behaviour. Perhaps application of the developed methodology to mutant strains with more subtle phenotypes can shed the light into these challenges in future work.

Multicellular self-organization is prevalent in biological systems but has proven challenging to study. There are complex feedback and compensatory mechanisms at the population level as well as pleiotropic effects of single mutations. Given significant heterogeneity of individual cell behaviors, small trends in behaviors between mutant strains could dissipate over time or in contrast could accumulate leading to differences in the emergent patterns. Our results demonstrate how careful quantification of cell behavior coupled to data-driven modeling approaches can predict these effects and pinpoint important synergies and compensatory mechanisms.

## 4 Materials and Methods

### 4.1 Bacterial Strains, Plasmids, and Growth Conditions

All *M. xanthus* strains were grown in CYE broth [1% Bacto casitone (Difco), 0.5% yeast extract (Difco), 10 mM 4-morpholinepropanesulfonic acid (MOPS) (pH 7.6), and 0.1% MgSO_4_] and development was induced on thin (10 ml in 100 mm Petri dish) TPM agar [10 mM Tris· HCl (pH 7.6), 1 mM KH(H_2_)PO_4_ (pH 7.6), 10 mM MgSO_4_, 1.5% agar (Difco)] plates containing 1 mM isopropyl *β*-D-1-thiogalactopyranoside (IPTG) and 100 *μ*M vanillate as described in [9]. Strain LS3910 was constructed by electroporation [38] of pLJS145 [9] into LS2442 [11] Transformants were selected using CYE 1.5% agar plates containing 15 *μ*g mL^−1^ oxytetracycline. *pilC* mutant LS3011 was constructed by Magellan mutagenesis of DK1622 as described in [39]. Strain LS4223 was constructed by electroporating the tdTomato plasmid pLJS145 into LS3011 with selection on CYE agar containing 15*μ*g mL^−1^ oxytetracycline.

### 4.2 Fluorescence Time-Lapse Microscopy

Time-lapse image capture was performed as described in [9]. As in [9], the beginning of aggregation varied between replicates by up to 1 h. To avoid possible bias in movie alignment caused by differences in the aggregation rate of WT and mutant cells, the approach of using the fraction of tdTomato cells within the aggregates used in [9] was replaced with a technique that relied on YFP fluorescence. To quantify aggregation progress using YFP fluorescence, the 2D Fourier transform coefficient magnitudes for wavelengths between 50 and 100 *μ*m were summed for each frame. Aggregation start was then detected as the point at which the summed magnitude in the movie frames crossed 20% of the maximum value reached in that movie. Movies were then cropped to align the detected beginning of aggregation and equalize their lengths as described in [9].

### 4.3 Developmental assays

The developmental assays were performed by mixing the tdTomato fluorescent strains LS4223, or LS3909 with the YFP fluorescent wild-type strain LS3630 in a 1:10,000 ratio. tdTomato fluorescence indicated positions of the individual cells for strains LS4223, and LS3909, while the YFP fluorescence revealed the territories of the LS3630 cell aggregates. Specifically, 100 L of the tdTomato-expressing strains and 1 mL of the YFP or non-fluorescent strains in the exponential phase were collected by centrifugation at 17, 000 × *g* for 1-2 min and washed with 100 L of ddH_2_O, respectively. The tdTomato strains were further diluted to 5 × 10^6^ cells mL^−1^, while the YFP or non-fluorescent strains were concentrated to 5 × 10^8^ cells mL^−1^ in ddH_2_O. The diluted tdTomato strains were then mixed with the YFP or non-fluorescent strains in a 1:100 ratio, resulting in a final ratio of 1:10,000 between the tdTomato and the YFP or non-fluorescent cells. 35 L of the cell mixtures in 4-6 replicates were spotted onto a TPM plate and dried out in a 32C° incubator for 30-45 min. The plate was sealed with parafilm and incubated in a 28C° dark room. With strains or mixtures that developed, development usually started between 7 and 10 hours post incubation and produced stable aggregates in another 5 to 8 hours. For time-lapse movies, images were captured at 30-sec intervals beginning about 1 hour before the initiation of aggregation and lasting until the formation of stable aggregates.

Viable spore data was obtained as noted previously [7].

### 4.4 Cell Tracking, Cell-State Detection, Run Vector Extraction, and Aggregate Tracking

Cell tracking, cell-state detection, and run vector extraction were performed as described in [9]. Time-lapse images were band-pass filtered to filter out the background fluorescent signal. Thereafter, the MATLAB function ’regionprops’ was used to identify the centroid and orientation of every fluorescently labeled cell. Image-to-image linking of detected cell positions into trajectories is achieved using the method introduced in ref. [17]. To detect movement characteristics of the cell, an extended Kalman filter (EKF) was developed to estimate the most likely movement state of the cell at a certain time-step. Here we assume cells will be in one of three different states: persistent forward, persistent backward, and non-persistent state. The EKF uses the anticipated cell position and detected cell position to calculate the probability of every state. The state with the maximum likelihood was then assigned as the movement state between the two images. Cell trajectories were then divided into run vectors, which start at the start of one contiguous movement state and end with the following change of state. The average speed, duration, distance, angle to the closest aggregate centroid, distance to the closest aggregate boundary were calculated for every run vector. The detection of aggregate is based on the light intensity of pixels. The threshold of the aggregate light intensity is calculated using K-means clustering on the pixels in the final frames of experiments. Areas with light intensity higher than the threshold are considered as aggregates. For simulations in fig. S6, this threshold is increased by 20%.

### 4.5 Data-Driven Agent-Based Model

The agent-based model used here is adapted from our previous work [9]. Given that simulations with the experimental mutant-to-WT ratio will lead to an unfeasible number of agents to simulate, we instead chose to implement the wide excess of WT cells via asymmetry in their interactions. We sample behaviors of both WT and mutant cells conditional only on the WT population distributions (see below). Each simulation consists of 10,000 WT agents and 8,000 mutant agents on a rectangular domain of 986 *μ*m× 740 *μ*m, equal to the microscope field of view, with periodic boundary conditions along each side. Each agent represents a single cell sampled from a biofilm of the same average density as in experiments (1.1 cells/*μ*m^2^), similar to sampling cell behaviors in the biofilm using a small number of fluorescently labeled cells. Similar to our previous model [9], each agent’s behaviors such as run speed, run duration, and run angle are drawn from the experiment data based on the time since the beginning of the experiment, the angle between the cell orientation and the average bearing angle of neighboring runs, and distance and angle to nearest aggregate. Note that unlike our previous model [9], here we did not use local cell density extracted from the time-lapse microscopy to choose our agent behaviors since the light intensity in the experiment varies too much for reliable density estimates outside the aggregates. We use the same method as in [9] to select run behavior for agents. Varying the number of mutant agent simulated does not significantly alter the results, e.g. fig. S5.

Since in the experiments the ratio of WT to mutant cells is over 10,000:1, it’s fair to assume that WT cell behavior is not affected by the mutant cells. Therefore, in the simulation, we usually chose the agent behavior based solely on the population distribution of WT agents. Moreover, the density estimation in simulation uses only WT agents so that mutant agents will not affect WT agent behavior. However, in simulations where we swap some WT data with mutant data, e.g., in fig. 8 and fig. 9, we do this by replacing some mutant data with WT data or vice versa, and feed the combined data to mutant agents. WT agents will always use WT data to provide background information such as neighbor cell alignment and density profile etc.

For simulations where agents use both mutant data and WT data (fig. 8 and fig. 9), the simulation process is slightly different from the original [9]. In particular, simulations where agents use WT data for non-persistent probability or non-persistent state behavior and mutant data for other behaviors, agents will choose their behaviors from WT data or mutant data accordingly using nearest-neighbor methods. A similar procedure is applied for simulations where agents use mutant data for non-persistent probability or non-persistent state behavior and WT data for other behaviors. For simulations where agents use WT data for persistent state speed or duration and mutant data for other behaviors, agents will choose their behaviors from mutant data only and then scale the persistent state speed or duration to match the mean of the WT data. This way, we can keep the correlation between the speed and duration in the mutant data. Moreover, to keep the bias in WT data, we split WT data into 2 branches: data of cells moving towards the aggregate and data of cells moving away from the aggregates. To keep the traffic-jam effect in WT data, we further split the 2 data branches into smaller branches based on the distance to aggregate: Each branch now contains data of cells with distance to aggregate within 1 *μ*m window and moving in the same direction. Then we calculate the mean speed or duration of each branch of data. We perform a similar calculation for mutant data and use the means to scale the agent behavior to match the WT data using the following equation:

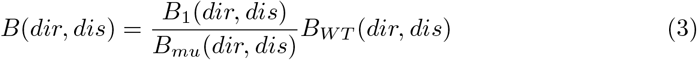

Where *B* is the final scaled behavior (speed or duration) for the agent, *dir* is the moving direction (moving towards or away from aggregate) of the agent and *dis* is the distance to aggregate, *B*_1_ is the selected behavior from mutant data, *B_mu_* the mean of the mutant data calculated as above and *B_WT_* is the mean of WT data calculated as above. For simulations where agents use mutant data for persistent state speed or duration and WT data for other behaviors, we apply a similar procedure, the equation to scale the agent behavior becomes:

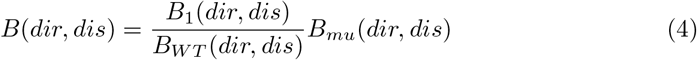

where *B*_1_ here is the selected behavior from WT data.

### 4.6 Data and Code Availability

The original tiff images recorded in the experiments and used for the analysis are available at datadryad.org (DOI:10.5061/dryad.1rn8pk0qc). All simulation and visualization codes are written in Matlab. The codes and resulting data for each figure are available at GitHub.

## 5 Acknowledgments

The research reported here was supported by the National Science Foundation DMS-1903275, IOS-1856742, MCB-1411891, and PHY-1427654 (for Center for Theoretical Biological Physics) and by the Welch Foundation (Grant C-1995).

## Contribution statement

Zhe Lyu and Christopher Cotter performed the experiments. Zhaoyang Zhang performed simulation and data analysis for experiments. Zhaoyang Zhang, Christopher Cotter, Lawrence Shimkets and Oleg Igoshin wrote the manuscript

## 6 Supplemental Material

**Figure S1.**
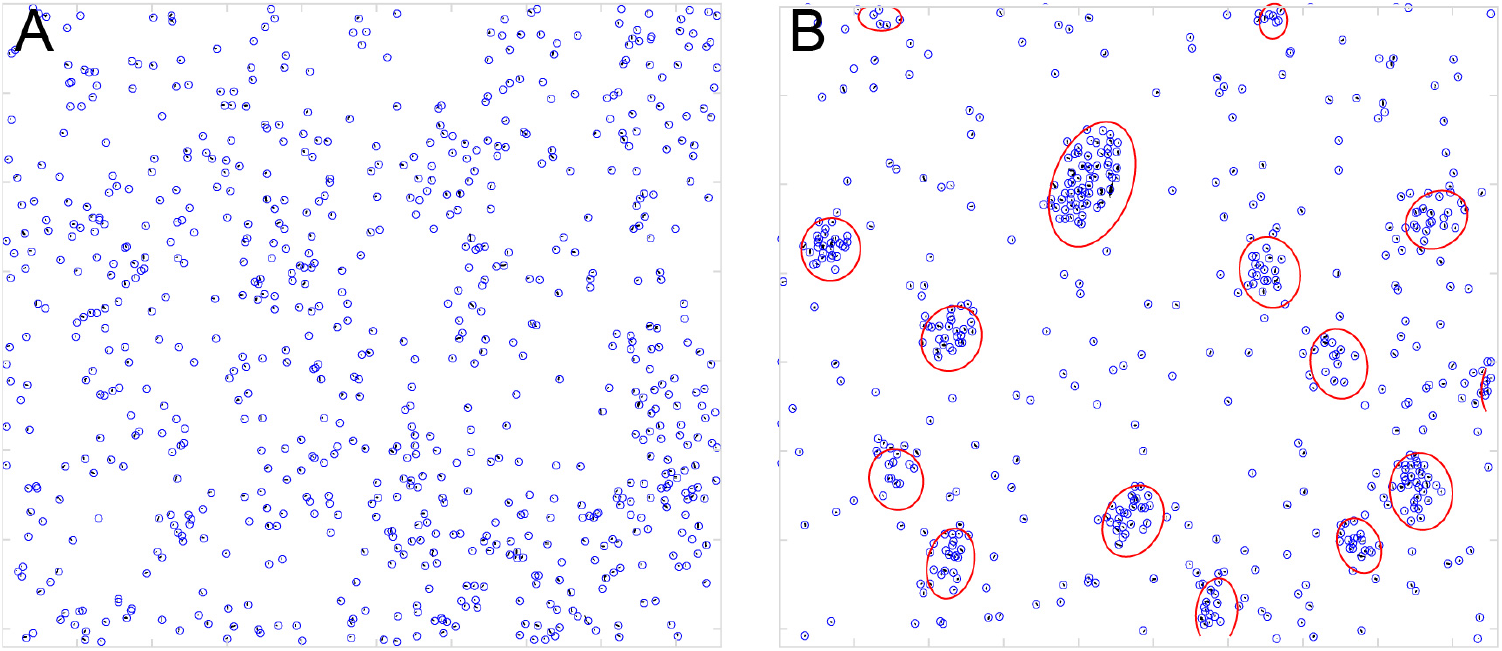
Aggregation result of WT cells from ref. [9]. (A): Beginning frame of WT experiment. (B): Ending frame of WT experiment. Blue circles are labeled cells, red circles are aggregates.

**Figure S2.**
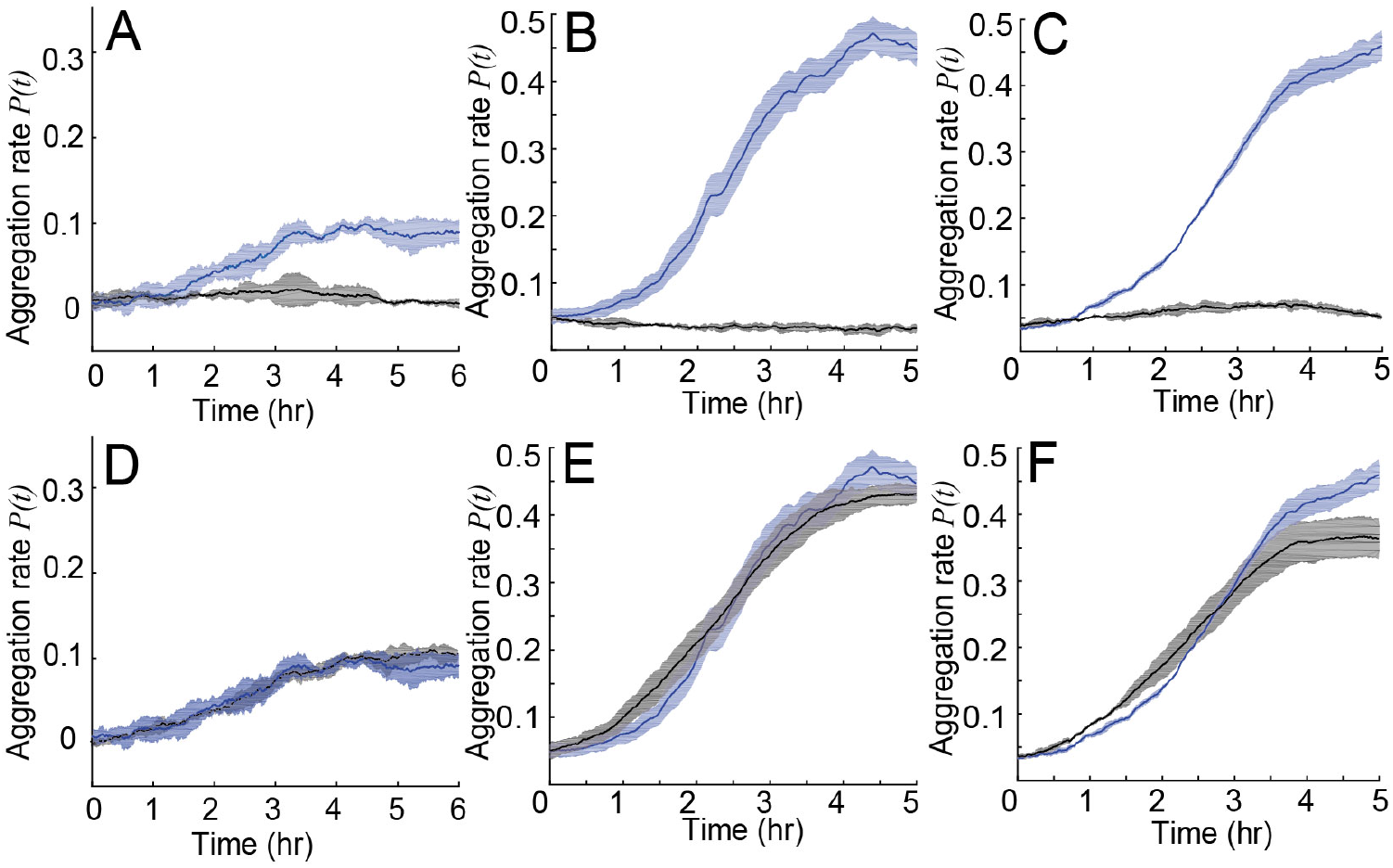
Identification of key cell behaviors that drive mutant strain aggregation. Simulation results of *pilC* (A,D), *csgA* (B,E) and WT (C,F) based on the experimental data (quantified as *P*(*t*), Eq. 1) on y-axis). Blue line and shaded areas are the simulation results under normal conditions. Black lines represent simulations without any dependence, i.e., data is randomly chosen (A-C) or simulation where run duration does not depend on time. (D-F). Shaded areas show standard deviations.

**Figure S3.**
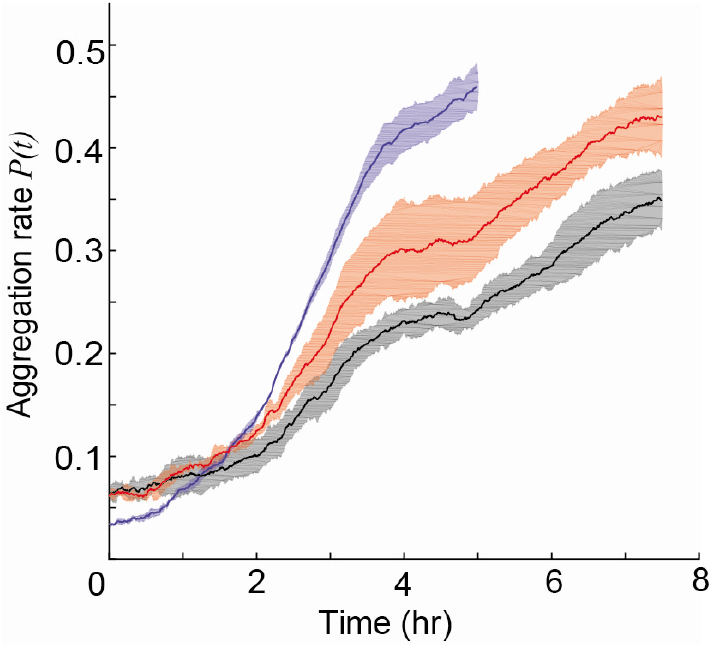
Simulation of WT agents with longer non-persistent duration (red) or higher non-persistent probability (black) impede aggregation rate (Eq. 1) as compared to simulations with unperturbed behaviors(blue). Shaded areas show standard deviations.

**Figure S4.**
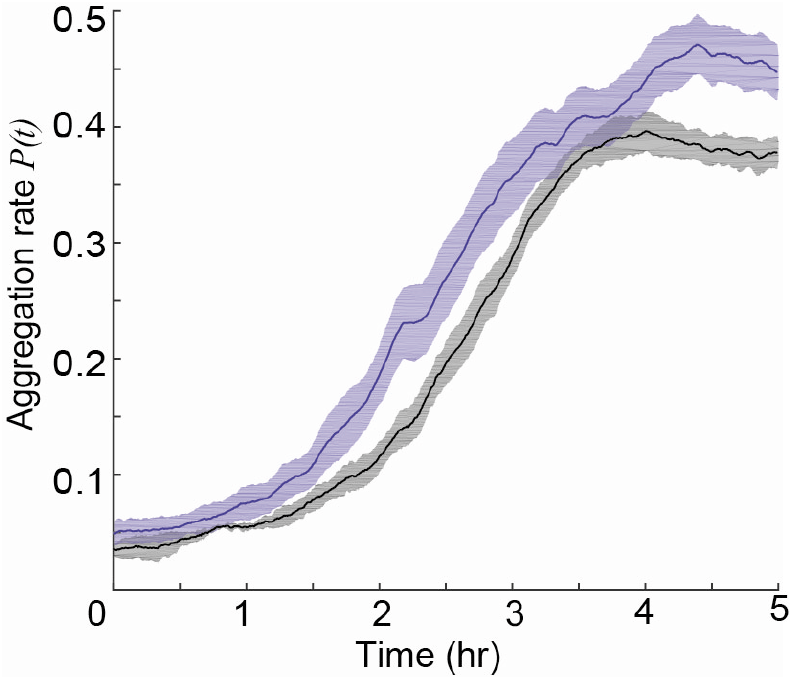
Model demonstrate that longer persistent run duration help *csgA*. Scaling persistent of *csgA* agents to WT persistent duration (black line) impedes their aggregation as compared to simulations using unscaled data(blue line). Shaded areas show standard deviations.

**Figure S5.**
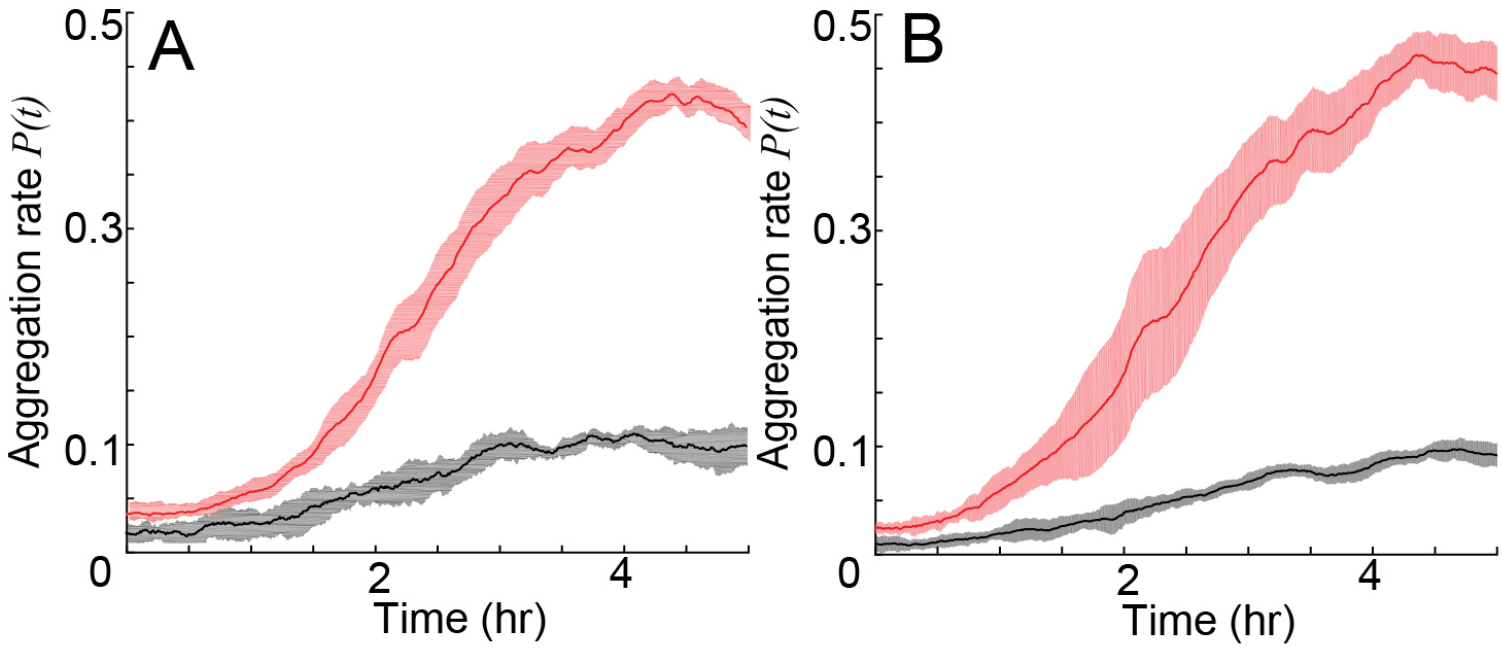
(A): Simulation result of mutant cells same as fig. 3. (B): Simulation result of mutant cells after increasing mutant agent to 10000. The results are similar, suggesting our agent number is enough.

**Figure S6.**
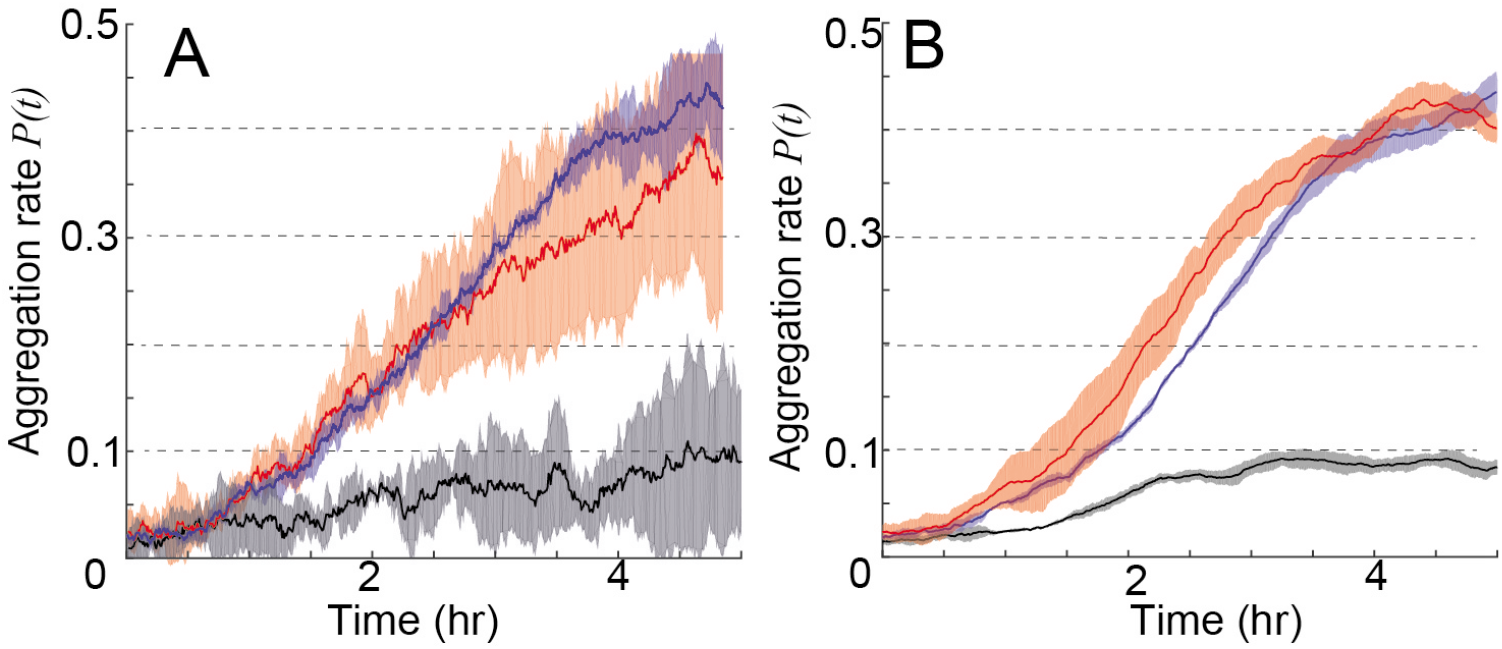
Result of aggregation rate after increasing the threshold for detecting aggregate in the experiment (A) and simulation (B). The results are very similar to fig. 3, suggesting that our results are robust regarding the threshold selection.

## Notes

### Competing Interest Statement

The authors have declared no competing interest.

